# Characterization and Transposon Mutagenesis of the Maize (*Zea mays*) *Pho1* Gene Family - Submission to PLOS Journals

**DOI:** 10.1101/040899

**Authors:** M. Nancy Salazar-Vidal, Edith Acosta-Segovia, Nidia Sanchez-Leon, Kevin R. Ahern, Thomas P. Brutnell, Ruairidh J. H. Sawers

## Abstract

Phosphorus is an essential nutrient for all plants, but also one of the least mobile, and consequently least available, in the soil. Plants have evolved a series of molecular, metabolic and developmental adaptations to increase the acquisition of phosphorus and to maximize the efficiency of use within the plant. In Arabidopsis (*Arabidopsis thaliana*), the AtPHO1 protein regulates and facilitates the distribution of phosphorus within the plant. To investigate the role of PHO1 in maize (*Zea mays*), the B73 reference genome was searched for homologous sequences and four genes identified that were designated *ZmPho1;1, ZmPho1;2a, ZmPho1;2b* and *ZmPho1;3. ZmPho1;2a* and *ZmPho1;2b* are the most similar to AtPho1, and represent candidate co-orthologs that we hypothesize to have been retained following whole genome duplication. Tissue-and phosphate-specific differences in the accumulation of *ZmPho1;2a* and *ZmPho1;2b* transcripts were observed, indicating regulatory divergence. Furthermore, evidence was obtained for the phosphate-regulated production of anti-sense transcripts associated with both *ZmPho1;2a* and *ZmPho1;2b*, suggesting the possibility of regulatory crosstalk between paralogs. To characterize functional divergence between *ZmPho1;2a* and *ZmPho1;2b*, a program of transposon mutagenesis was initiated using the Ac/Ds system, and, here, we report the generation of novel alleles of *ZmPho1;2a* and *ZmPho1;2b*.

## Introduction

Phosphorus (P) is an essential nutrient for all plants and a limitation on productivity in many agricultural systems [1]. Current levels of agricultural phosphorus inputs are recognized to be both unsustainable and environmentally undesirable [2]. Rational strategies to improve P efficiency in agricultural systems demand a greater understanding of P relations in crop plants, both in terms of P uptake from the soil and P translocation and use within the plant.

The protein PHO1 has been characterized in Arabidopsis (*Arabidopsis thaliana*) and rice (*Oryza sativa*) to play a key role both in the export of inorganic P (Pi) to the xylem apoplast for translocation [3] and in the modulation of long-distance signals underlying the P-deficiency response [4]. The Arabidopsis *Atpho1* mutant hypo-accumulates P in the shoots and displays associated symptoms of phosphate deficiency, including reduced growth rate, thinner stalks, smaller leaves, very few secondary inflorescences, delayed flowering and elevated levels of anthocyanin accumulation [3]. In rice, disruption of *OsPHO1;2*, the ortholog of *AtPHO1*, results in a phenotype similar to that of the *Atpho1* mutant [5], suggesting that the two genes are functionally equivalent. Indeed, expression of *OsPHO1;2* in the *Atpho1* background was found to partially complement the mutant phenotype [6]. A feature distinguishing the rice *OsPHO1;2* gene from its Arabidopsis ortholog is the P-regulated production of a *cis*-Natural Antisense Transcript (*cis*-NAT _OsPHO1;2_) [5] [7] which acts as a translational enhancer [8].

PHO1 proteins contain two conserved domains: the N-terminal hydrophilic SPX domain (named for the yeast proteins Syg1 and Pho81, and the human Xpr1) and the C-terminal hydrophobic EXS domain (named for the yeast proteins ERD1 and Syg1 and the mammalian Xpr1) [9]. The SPX domain is subdivided into three well-conserved sub-domains, separated from each other by regions of low conservation. SPX domain containing proteins are key players in a number of processes involved in P homeostasis, such as fine tuning of Pi transport and signaling by physical interactions with other proteins [7]. Following the SPX domain there are a series of putative membrane-spanning α-helices that extend into the C-terminal EXS domain [9]. In AtPHO1, the EXS domain is crucial for protein localization to the Golgi/*trans*-Golgi network and for Pi export activity as well as playing a role in the modulation of long-distance root-to-shoot signaling under P limitation [4].

Despite the importance of maize as a staple crop and the dependence of maize production on large-scale input of phosphate fertilizers, the molecular components of maize P uptake and translocation remain poorly characterized [10]. Although it has been possible to identify maize sequences homologous to known P-related genes from other species, functional assignment has been based largely on patterns of transcript accumulation. With the development of accessible public-sector resources, it is now feasible to conduct reverse genetic analyses in maize. Here, we extend the molecular characterization of maize P response by generating mutant alleles of maize *Pho1* genes using endogenous *Activator/Dissociation (Ac/Ds)* transposable elements. The *(Ac/Ds)* system consists of autonomous *Ac* elements that encode a transposase (TPase) and non-autonomous *Ds* elements that are typically derived from *Ac* elements by mutations within the TPase gene. Lacking TPase, *Ds* elements are stable, unless mobilized by TPase supplied in trans by an *Ac. Ac/Ds* elements move via a cut-and-paste mechanism [11], with a preference for transposition to linked sites [12] that makes the system ideal for local mutagenesis [13]. To exploit the system for reverse genetics, *Ac* and *Ds* elements have been distributed throughout the genome and placed on the maize physical map, providing potential “launch pads” for mutagenesis of nearby genes [14] [15].

In this study, we identify four maize *Pho1* like genes in the maize (var. B73) genome, including two (*ZmPho1;2a* and *ZmPho1;2b*) that we consider co-orthologs of *AtPHO1.* Structures of the *ZmPho1;2a* and *ZmPho1;2b* genes was experimentally determined and accumulation of transcripts was characterized in the roots and shoots of seedlings grown in P-replete or P-limiting conditions. Novel insertional alleles of *ZmPho1;2a* and *ZmPho1;2b* are reported, generated using the *Ac/Ds* transposon system. Availability of mutant alleles will be central in determining the functional role of *ZmPho1;2a* and *ZmPho1;2b* in maize.

## Materials and Methods

### Identification of maize *Pho1* genes

The AtPHO1 cDNA sequence (GenBank ID: AF474076.1) was used to search the maize working gene set peptide database (www.maizesequence.org) in a BLASTX search performed under the default parameters. Identified maize sequences were in turn used to reciprocally search *Arabidopsis thaliana* (www.phytozome.net). Four sequences were identified with a high level of similarity to AtPHO1: GRMZM5G891944 (chr 3:28,919,073-28,923,871); GRMZM2G466545 (chr 4:171,946,555-171,952,268); GRMZM2G058444 (chr 5:215,110,603-215,115,635) and GRMZM2G064657 (chr 6:122,577,593-122,582,074). The putative protein sequences were confirmed to contain canonical PHO1 domain structure by PFam analysis (pfam.sanger.ac.uk) and NCBI conserved domains search (www.ncbi.nlm.nih.gov).

### Amplification of full length *Pho1;2* cDNAs

Total RNA was extracted using Trizol-chloroform from the roots of 10-day-old B73 seedlings grown under phosphate limiting conditions (sand substrate; fertilized with 0mM PO4). 1.5μg of RNA was used to synthesize cDNA with oligo(dT) primer and *SuperScript III* Reverse Transcriptase (Invitrogen, Carlsbad, CA, USA) in a reaction volume of *20*μl. PCR amplification of full length *Pho1;2* cDNAs was performed using using Platinum Taq DNA Polymerase High Fidelity (Invitrogen) under the following cycling conditions: initial incubation at 95°C for 3min, followed by 35 cycles of 94°C for 30sec, 61°C for 30sec and 68°C for 30sec, final extension at 68°C for 5min. Primers used are shown in Table 1.

**Table 1.**
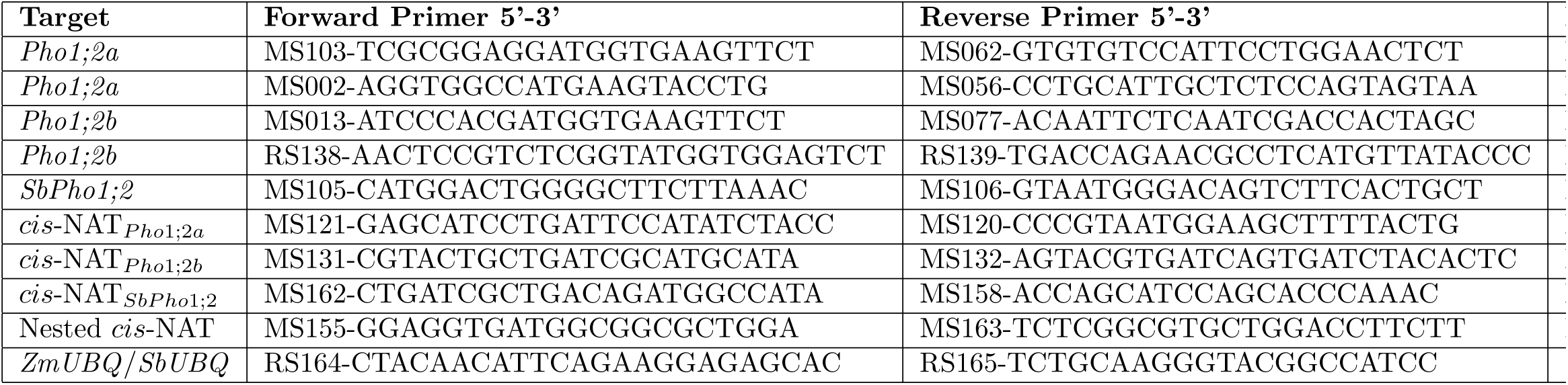
Primers used in this study for CDS amplification and transcript accumulation

### Analysis of *Pho1;2* transcript accumulation

Total RNA was extracted using Trizol-Chloroform from the roots of 10-day-old B73 seedlings grown under P replete (sand substrate; fertilized with 1mM PO4) or P limiting (sand substrate; fertilized with 0mM PO4) conditions. cDNA was synthesized as described above. PCR amplification of sense genes was performed with the primers MS002-MS056 *(Pho1;2a)*, RS138-RS139 *(Pho1;2b)* and MS105-MS106 *(SbPho1;2)* under the following cycling conditions for: initial incubation at 95°C for 5min, followed by 32 cycles of 95°C for 30sec, 63°C for 30sec and 72°C for 1min, final extension 72°C for 5min. PCR amplification of *cis*-NATs was performed with the primers MS121-MS120 (*cis*-NATP_*Pho1;2a*_), MS131-MS132 (*cis*-NATP_*Pho1;2b*_) and MS162-MS158 (*cis*-NATSbPho1_*SbPhol;2*_) under the following cycling conditions for: initial incubation at 95°C for 5min, followed by 38 cycles of 95°C for 30sec, 59°C for 30sec and 72°C for 30sec, final extension 72°C for 5min. PCR products from this first PCR reactions were diluted 1:100,000 and used as templates for nested PCR with MS155-MS163 primers for all products and following conditions: initial incubation at 95°C for 5min, followed by 25 cycles of 95°C for 30sec, 58°C for 30sec and 72°C for 15sec, final extension 72°C for 5min. Primers are shown in Table 1. Maize and sorghum poly-ubiquitin (GRMZM2G419891/Sb04g004260) were used as a control, along with amplification from genomic DNA template, using 50ng gDNA in *20*μl for 32-cycle reactions and 10ng gDNA in *20*μl for 38-cycle reactions. Products were analyzed on 1.5% agarose gels.

### Transposon mutagenesis

The strategy for *Ac/Ds* mutagenesis was as previously described [16] [14]; [15]. Genetic stocks were maintained in the T43 background, a color-converted W22 stock carrying *r1-sc∷m3*, a Ds6-like insertion in the *r1* locus that controls anthocyanin accumulation in aleurone and scutellar tissues [17]. The frequency of purple spotting in the aluerone resulting from somatic reversion of *r1-sc∷m3* was used to monitor Ac activity [18]. Donor Ac and Ds stocks were selected from existing collections [14] [15]: the element *bti31094∷Ac* is placed on the B73 physical map 650.8Kb from *ZmPho1;2a*; the element *I.S06.1616∷Ds* is inserted in intron 13 of *ZmPho1;2b*, and was subsequently designated *ZmPho1;2b-m1∷Ds*. To generate a testcross population for mutagenesis of *ZmPho1;2a*, 207 individuals homozygous for *bti31094∷Ac* were crossed as females by T43, and rare, finely spotted progeny kernels were selected for screening. To re-mobilize the Ds element *I.S06.1616∷Ds* within *ZmPho1;2b*-1, homozygous *ZmPho1;2b∷Ds* individuals carrying the unlinked stable transposase source Ac-Immobilized Ac-im [19], were used as males to pollinate T43, and coarsely spotted progeny kernels were selected for screening.

To identify novel *Ac/Ds* insertions in *pho1;2a* and *pho1;2b*, selected kernels were germinated in the greenhouse and DNA was isolated from pools of 18 seedlings. The candidate gene space was explored by PCR using a range of gene-specific primers in combination with “outward-facing” *Ac/Ds*-end primers. All primers are listed in Table 2. PCR reactions contained 400ng gDNA and .25μM of each primer. Reactions for *pho1;2a* were performed using Platinum Taq DNA Polymerase High Fidelity (Invitrogen, Carlsbad, CA, USA) under the following cycling conditions: denaturation at 94°c for 4 min; 30 cycles of 94°c for 30 sec, 58°c for 30 sec, 68°c for 3min 30 sec; final extension at 68°̱c for 10 min. Reactions for *pho1;2b* were performed using Kapa Taq DNA polymerase (Kapa Biosystems, Wilmington, Massachusetts, USA) under the following cycling conditions: denaturation at 95°c for 5 min; 35 cycles of 95°c for 30 sec, 58°c for 30 sec, 72°c for 3min 30 secs; final extension at 72°c for 5 min. Positive pools were re-analyzed as individuals, following the same cycling conditions. PCR reactions were analyzed on 1.5% agarose gels. Products from positive individuals were purified using QIAquick PCR Purification Kit (Qiagen, Hilden, Germany), ligated into pGEM T-easy vector (Promega, Fitchburg, Wisconsin, USA) and sequenced. Genotyping of *ZmPho1;2a-m1∷Ac* was performed by amplification of flanking insertion sides with specific-Ac primers (MS124-MS052, MS124-JRS01 and MS052-JGp3) shown in Table 2 with Kapa Taq DNA polymerase (Kapa Biosystems), under the following cycling conditions: denaturation at 95°c for 5 min; 35 cycles of 95°c for 30 sec, 58°c for 30 sec, 72°c for 3min 30 secs; final extension at 72°c for 5 min, and confirmed by CAPS assays using MS124-MS052 PCR product and BseYI restriction enzyme (New England BioLabs, Ipswich, Massachusetts, USA) protocol.

**Table 2.**
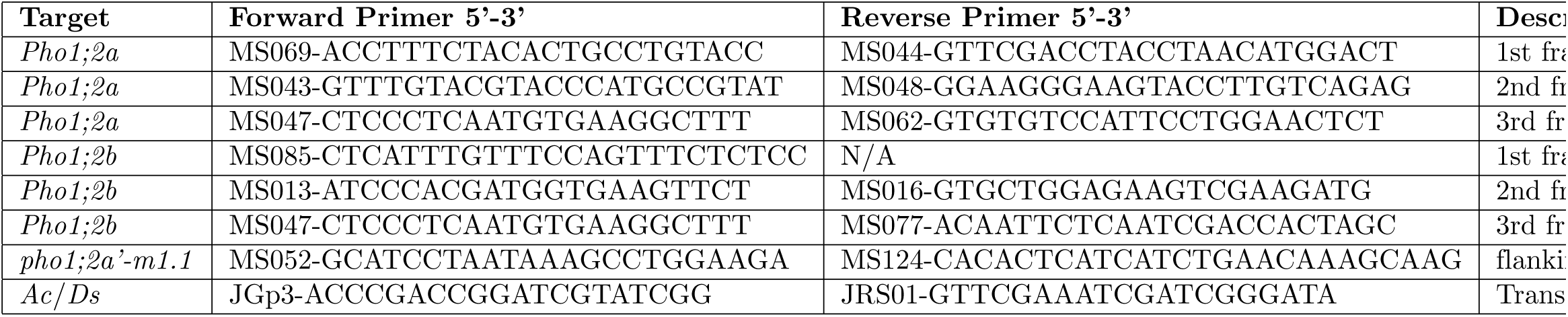
Primers used in this study for genotyping

To generate footprint alleles, individuals homozygous for *pho1;2a-m1∷Ac* were crossed as males to T43 females. Rare, non-spotted progeny kernels were selected and screened for excision by PCR amplification across the site of Ac insertion using primers MS124-MS052 shown in Table 2 with Kapa Taq DNA polymerase (Kapa Biosystems), under the following cycling conditions: denaturation at 95°c for 5 min; 35 cycles of 95°c for 30 sec, 58°c for 30 sec, 72°c for 3min 30 secs; final extension at 72°c for 5 min. PCR products from each individual were purified using QIAquick PCR Purification Kit (Qiagen), ligated into pGEM T-easy vector (Promega) and sequenced. Genotyping was performed by CAPS assays using previous PCR protocol and BseYI restriction enzyme (New England BioLabs, Ipswich, Massachusetts, USA) protocol.

## Results

### The maize genome contains four PHO1 homologs

To identify maize *Pho1* genes, the B73 reference genome (B73 RefGen v3; www.maizegdb.org) was searched to identify gene models whose putative protein products exhibit a high degree of similarity to the *Arabidopsis* protein AtPHO1 (Table 3). Four such maize gene models were identified, and, on the basis of similarity to previously annotated rice genes [5], designated *ZmPho1;1* (GRMZM5G891944), *ZmPho1;2a* (GRMZM2G466545), *ZmPho1;2b* (GRMZM2G058444) and *ZmPho1;3* (GRMZM2G064657). To investigate orthology among *Arabidopsis thaliana* and *grass PHO1* genes, additional sequences were identified from sorghum (*Sorghum bicolor*) and canola (*Brassica rapa*), and used to generate a multiple alignment and distance tree (Fig. 1).

**Figure 1.**
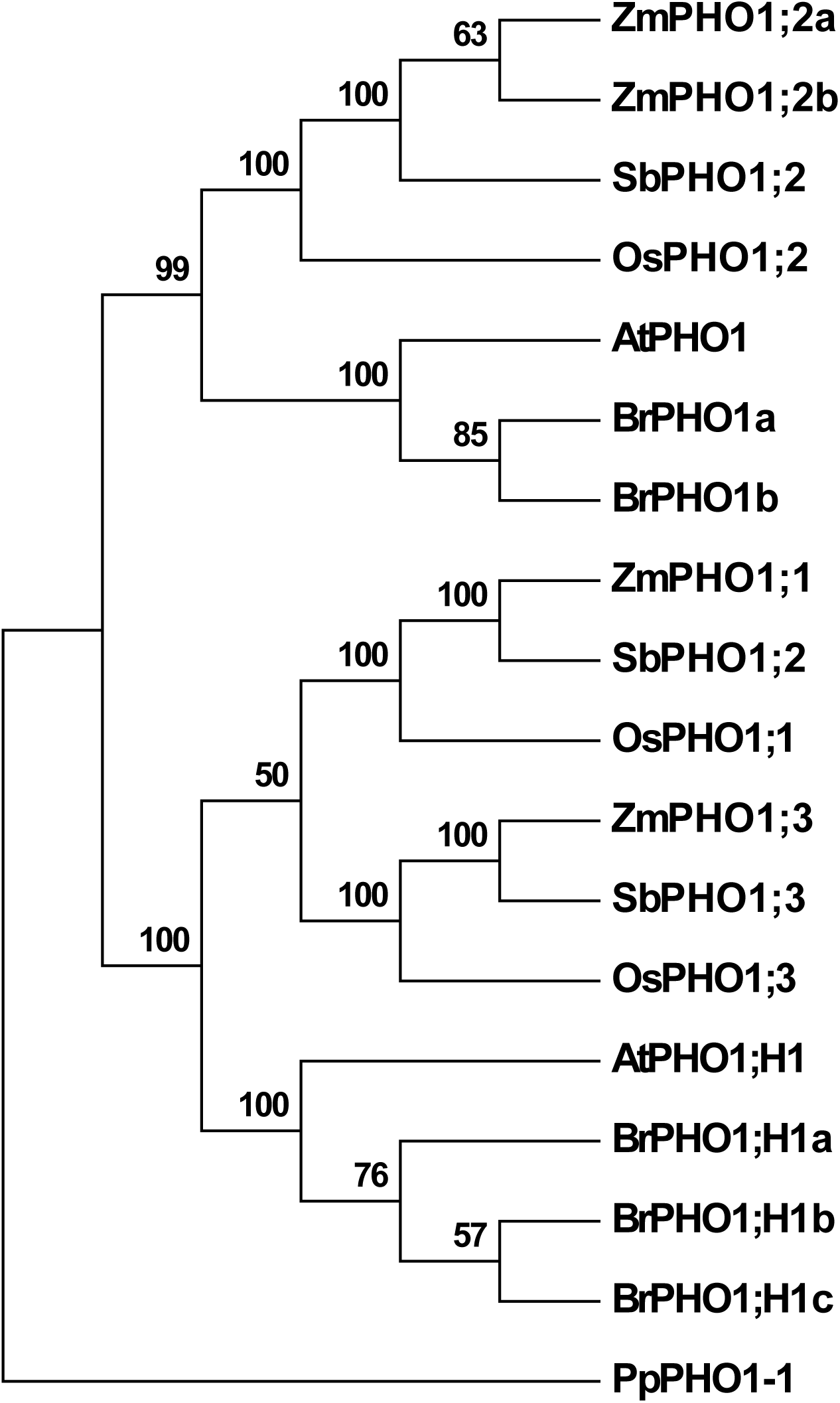
Maximum likelihood reconstruction of the complete set of maize (Zm), sorghum (Sb) and rice (Os) PHO1 proteins along with selected PHO1 proteins of Arabidopsis (At) and canola (Br). The moss (*Physcomitrella patens* protein PpPHO1-1 is shown as an outgroup. Bootstrap support shown as a percentage at the nodes.

**Table 3.** Maize Pho1 genes and corresponding *Arabidopsis* and rice orthologs

| *At* Gene | *Os* Gene | *Zm* Gene | *Zm* Gene Model | *Zm* Position |
| --- | --- | --- | --- | --- |
| *AtPHO1* | *OsPho1;2* | *ZmPho1;2a* | GRMZM2G466545 | chr 4:171,946,555-171,952,268 |
| *AtPHO1* | *OsPho1;2* | *ZmPho1;2b* | GRMZM2G058444 | chr 5:215,110,603-215,115,635 |
| *AtPHO1;H1* | *OsPho1;1* | *ZmPho1;1* | GRMZM5G891944 | chr 3:28,919,073-28,923,871) |
| *AtPHO1;H1* | *OsPho1;3* | *ZmPho1;3* | GRMZM2G064657 | chr 6:122,577,593-122,582,074 |

From *Arabidopsis*, only the proteins AtPHO1 and AtPHO1;H1 were included in the analysis, leaving aside a large clade of divergent functionally distinct PHO1 proteins that are specific to dicotyledonous plants [20]. The analysis supported the previously reported divergence of PHO1 and PHO1;H1 clades, dating from before the divergence of monocotyledonous and dicotyledonous plants [5,20]. Within the PHO1;H1 clade, a duplication event was observed specific to the grasses in the analysis. As a result, the three grass species each contain two co-orthologs of AtPHO1;H1 – encoded by the genes annotated *Pho1;1* and *Pho1;3*. We observed also an expansion of the PHO1;H1 clade in canola, although this expansion is lineage specific, and there is no indication that Pho1;H1 was not a single gene at the base of this clade. The PHO1 clade itself contains the products of single-copy *Pho1 /Pho1;2* sequences in all species in our analysis, with the exception of a lineage-specific duplication in maize. As a consequence, the paralogous maize genes *ZmPho1;2a* and *ZmPho1;2b* are considered to be co-orthologos to *AtPHO1*.

### *ZmPho1;2a* and *ZmPho1;2b* show features of syntenic paralogs retained following to genome duplication

The high degree of sequence similarity between *ZmPho1;2a* and *ZmPho1;2b* suggests that they result from a recent gene duplication event. It has been hypothesized that the last whole genome duplication (WGD) event in maize occurred between 5 and 12 million-years-ago, sometime after divergence from the sorghum lineage, as the result of polyploidization [21]. The observation that *Pho1;2* is a single copy sequence in both rice and sorghum is consistent with the maize duplication arising during this most recent WGD. Further inspection revealed that the two genomic regions carrying maize *Pho1;2* genes are both syntenic to the sorghum region carrying *SbPho1;2*, and that the two maize regions have been assigned previously o distinct pre-tetraploid ancestral genomes (Chr4:168,085,162..179,711,318 to sub-genome 2; Chr5:208,925,180..217,680,842 to sub-genome 1; [21]). The return of the maize genome to a diploid organization following WGD has been accompanied by the loss of the majority of duplicate genes through a process known as fractionation [21]. In certain cases, however, pairs of syntenic paralogs have been retained. The genomic region surrounding the Pho1;2 genes exhibits a number of such candidate pairs in addition to *Pho1;2a* and *Pho1;2b* (Fig. 2A), providing ample evidence of micro-synteny between the regions. In both sorghum and maize *Pho1;2* genes are adjacent to a putative WD40 protein encoding gene, present on the opposite strand and partially overlapping the annotated 3’ UTR region of the *Pho1;2* sequence (Fig. 2B), a feature not observed in the other maize or sorghum *Pho1* paralogs.

**Figure 2.**
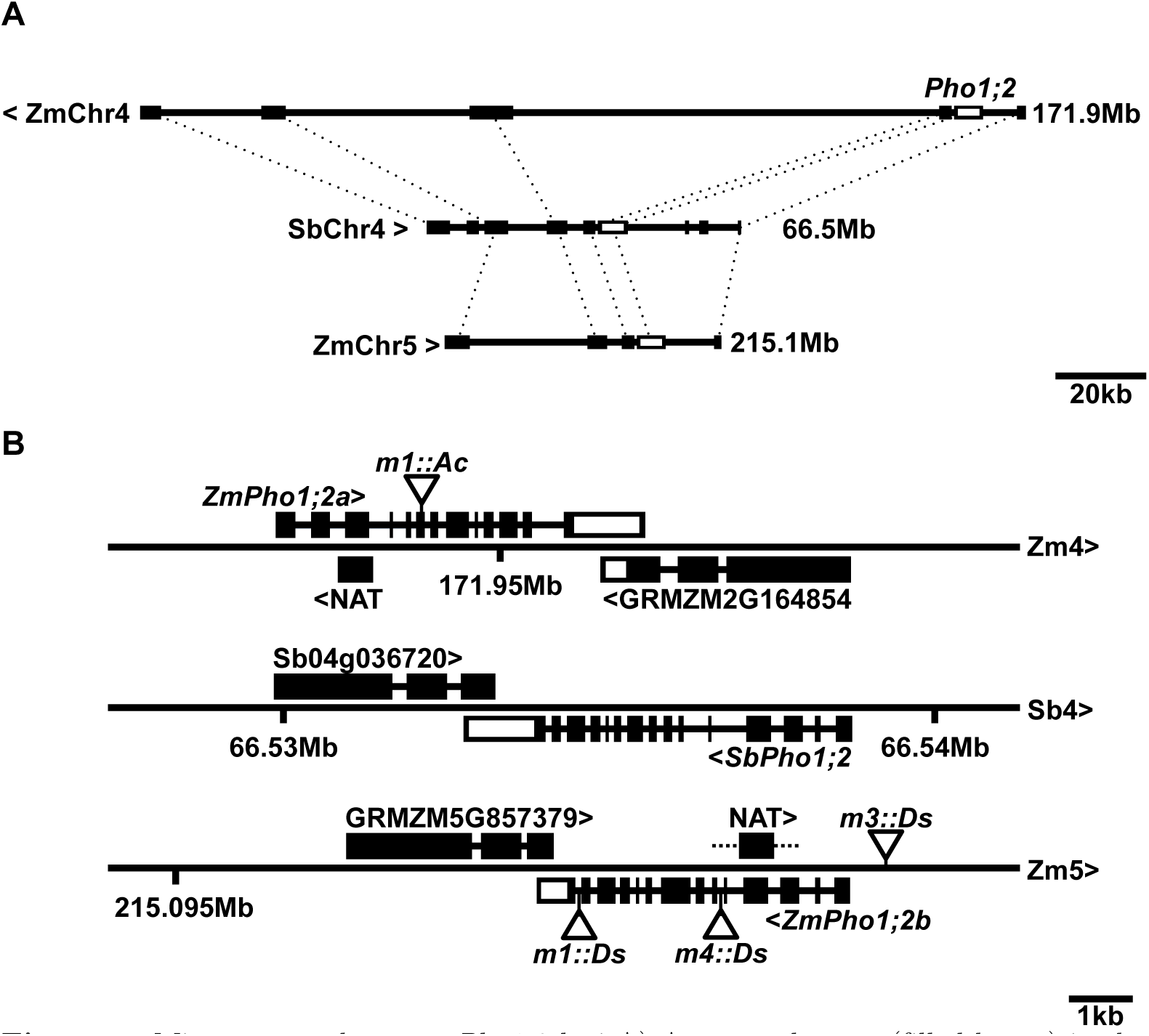
Microsynteny among *Pho1;2* loci A) Annotated genes (filled boxes) in the region of *SbPho1;2* on Sorghum chromosome 4 (SbChr4) and candidate orthologs on maize (B73) chromosomes 4 (ZmChr4) and 5 (ZmChr5). Orthologous genes are connected by dashed lines. *Pho1;2* genes on the three chromosomes shown in unfilled boxes. Regions shown to scale, the right hand position indicated. SbChr4 and ZmChr5 run left to right, ZmChr4 right to left. B) *Pho1;2* gene models on ZmChr4, SbChr4 and ZmChr5. Exons are shown as boxes, coding regions filled, UTR unfilled. Putative anti-sense transcripts associated with *ZmPho1;2a* and *ZmPho1;2b* shown as filled boxes (NAT). Angle brackets indicate the direction of transcription. The maize syntenic paralog pair GRMZM2G164854/GRMZM5G853379 and their sorghum ortholog are also shown. Triangles indicate the position of *Activator* and *Dissociation* insertion. Refer to Table 4 for full description alleles. Shown to scale.

### Transcripts encoded by *ZmPho1;2a* and *ZmPho1;2b* exhibit divergent patterns of expression

To determine the pattern of accumulation of transcripts encoded by *ZmPho1;2a* and *ZmPho1;2b*, RT-PCR was used to amplify gene-specific fragments of the two genes from cDNA prepared from roots or leaves of 10-day-old seedlings (B73) grown in sand, watered with either complete (+P) or a modified phosphate-free Hoagland solution (-P) (Fig. 3A). Transcripts of *ZmPho1;2a* were detected in roots but not leaves, with an indication of greater accumulation under -P. In contrast, transcripts of *ZmPho1;2b* showed constitutive accumulation in both roots and leaves, indicating regulatory divergence between the two maize *Pho1;2*. The accumulation of *SbPho1;2* transcripts in sorghum (BTx623) seedlings was also examined, under the same growth conditions. Transcripts of *SbPho1;2* accumulated in a pattern similar to that observed for *ZmPho1;2b*, suggesting that constitutive expression was the ancestral, pre-WGD, state. Subsequently, additional gene-specific PCR primers were designed and used to amplify the complete *ZmPho1;2a* and *ZmPho1;2b* cDNAs that were sequenced to confirm the gene-model structure.

**Figure 3.**
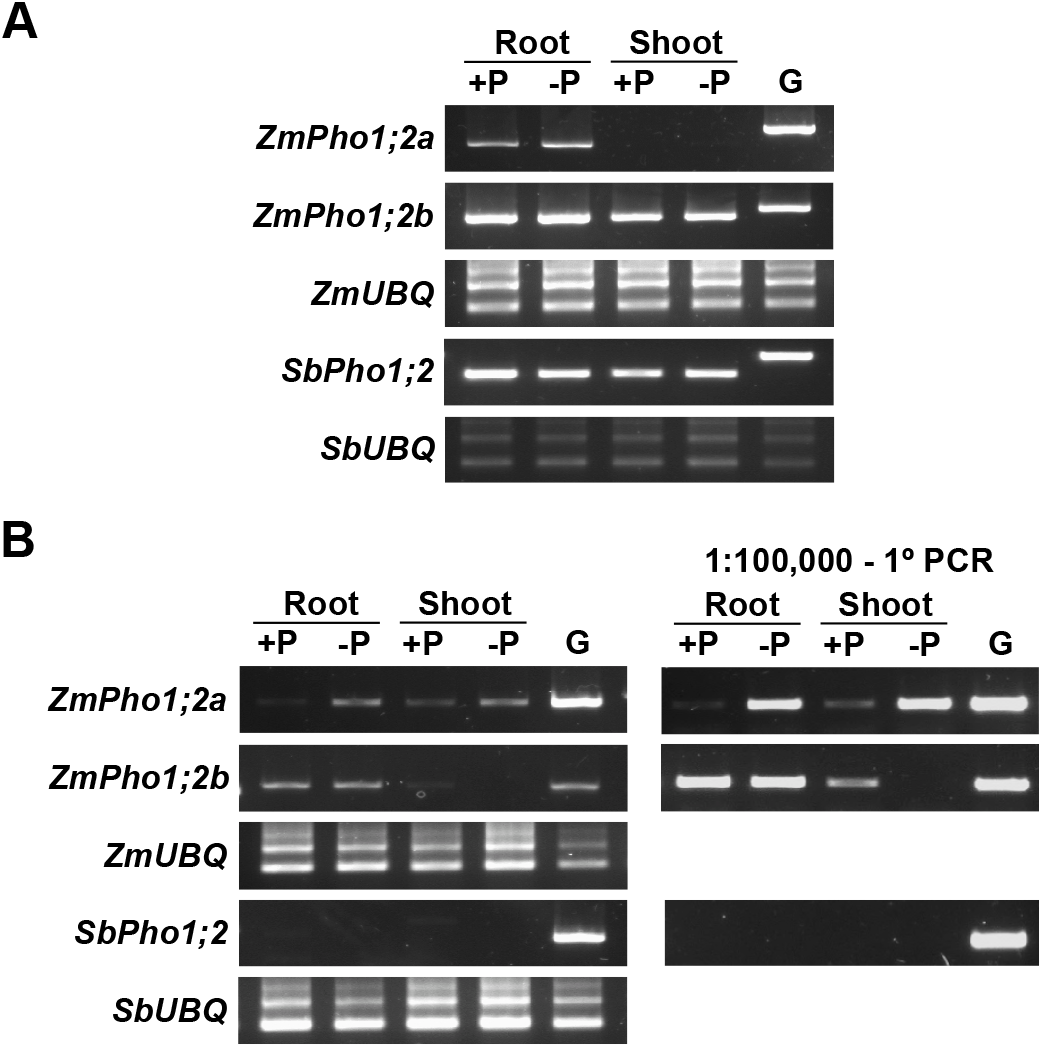
Accumulation of *Pho1;2* sense and antisense transcripts under contrasting phosphate conditions. (A) Amplification of fragments corresponding to the mature sense-RNAs of *ZmPho1;2a, ZmPho1;2b* and *SbPho1;2* from oligo-dT primed cDNA prepared from roots and shoot of 10-day-old B73 seedlings, fertilized with Hoagland solution adjusted to 1mM (+P) or 0mM (-P) inorganic phosphate. (B) Amplification of fragments corresponding to putative anti-sense transcripts encoded by *ZmPho1;2a, ZmPho1;2b* from cDNA as Panel A. Primary PCR (left) and nested PCR performed from 1:100,000 dilution of primary reaction (right). Amplification of *ZmUBQ* and *SbUBQ* fragments and amplification from genomic DNA template (G) were used as controls.

### *ZmPho1;2a* and *ZmPho1;2b* are associated with phosphate-regulated putative *cis*-natural anti-sense transcripts

Although we observed differential accumulation of ZmPho1;2 transcripts with respect to P availability, it has been reported that the Pho1;2 in rice is largely regulated at the post-transcriptional level by a P-regulated *cis*-Natural Anti-sense Transcript (*cis*-NAT_*OsPho1;2*_) [5]; [8]. The (*cis*-NAT_*OsPho1;2*_) transcript has been shown to act as a translational enhancer, and has been proposed to act by direct interaction with the sense transcript [8]. The rice *cis*-NAT_*OsPho1;2*_ initiates in Intron 4 of OsPho1;2 and extends into the 5’ UTR region [8]. A putative *ZmPho1;2a* anti-sense sequence is annotated in the maize reference genome in a homologous position to the rice transcript (www.maizegdb.org), although, on the basis of cDNA evidence, the transcript is considerably shorter than *cis*-NAT_*OsPho1;2*_, being trunctated at the 3’ end and extending only as far as Intron 2 of *ZmPho1;2a* (Fig. 2B). No paralogous sequence has been annotated associated with *ZmPho1;2b*.

To investigate the presence of *cis*-NAT transcripts associated with *ZmPho1;2a* and explore the possibility that a paralogous *cis*-NAT transcript might be generated from *ZmPho1;2b*, gene-specific primers were designed to the introns flanking the homologous Exons 4 and 3 of *ZmPho1;2a* and *ZmPho1;2b*, respectively. These primers were then used to attempt to amplify products from cDNA prepared from seedling root and leaves as described above. Products of the predicted size were successfully amplified using both *ZmPho1;2a* and *ZmPho1;2b* primer sets, consistent with the accumulation of *cis*-NATs (Fig. 3B). No products were amplified from no-RT control samples (data not shown). Putative *cis*-NAT products were sequenced and confirmed to originate from the *ZmPho1;2a* and *ZmPho1;2b* genes. There was no evidence of the accumulation of alternatively or partially spliced transcripts during the previous amplification of full length Pho1;2 cDNAs.

The accumulation of the putative *cis*-NAT_*ZmPho1;2a*_ was observed to be induced under-P conditions in both roots and leaves. In contrast, the putative *cis*-NAT_*ZmPho1;2b*_ transcript was observed to accumulate in roots and leaves, with no response to P availability (Fig.3B), providing further evidence of functional divergence between the paralogs. Interestingly, using the approach we employed in maize, we found no evidence of an equivalent *cis*-NAT associated with *SbPho1;2* (Fig. 3B), although additional experiments will be required to rule out the possibility that anti-sense transcripts are produced from other regions of the sorghum gene.

### Transposon mutagenesis of *ZmPho1;2a* and *ZmPho1;2b*

To investigate functional divergence between *ZmPho1;2a* and *ZmPho1;2b*, we initiated a program to mutagenize both loci using the endogenous *Activator/Dissociation (Ac/Ds)* transposon system. Ac and Ds elements show a strong preference for linked transposition, allowing a given element to be used for mutagenesis of nearby candidate genes. Once established, it becomes possible to generate multiple alleles from a single test-cross population.

To mutagenize *ZmPho1;2a*, we recovered 1082 novel transposition events from the element Ac (*Bti31094∷Ac*) located 650.8kb upstream the target (Fig. 4) by selection of rare high Ac dosage kernels (Fig.4) from a testcross population. A PCR-based strategy was designed to screen for reinsertion of Ac into *ZmPho1;2a*. The gene was divided into three overlapping fragments, and, allowing for both possible orientations of Ac insertion, we performed a total of 12 reactions to cover the gene space, screening first pools of 18 seedlings, and subsequently the individuals constituting positive pools. Putative insertions were re-amplified using DNA extracted from a second seedling leaf to reduce the probability of selecting somatic events. Using this strategy, we recovered a novel germinal Ac insertion in Exon 6 of *ZmPho1;2a* (*Zmpho1;2a-m1∷Ac*) (Fig.5). Left and right flanking border fragments were amplified and sequenced, confirming the exact location of the element and identifying an 8bp target site duplication (AGCCCAGG) consistent with *Ac* insertion. Further analysis of progeny recovered from original the positive plant revealed a high-frequency of somatic excision of the *Ac* element: apparently wild-type fragments were routinely amplified from individuals selected on the basis of kernal spotting pattern to be homozygous for *Zmpho1;2a-m1∷Ac*. Sequencing of these products revealed them to be the products of somatic *Ac* excision, as indicated by a short, typically 8bp, duplication adjacent to the former *Ac* insertion site. Excision of *Ac* from *Zmpho1;2a-m1∷Ac* leaving 7 or 8 bp of duplicated sequence results in the generation of a *BseYI* restriction site (CCCAGC) providing a means to demonstrate further *Ac* excission by direct digestion of PCR products (Fig. 6A,B,D) (Table 4).

**Table 4.**
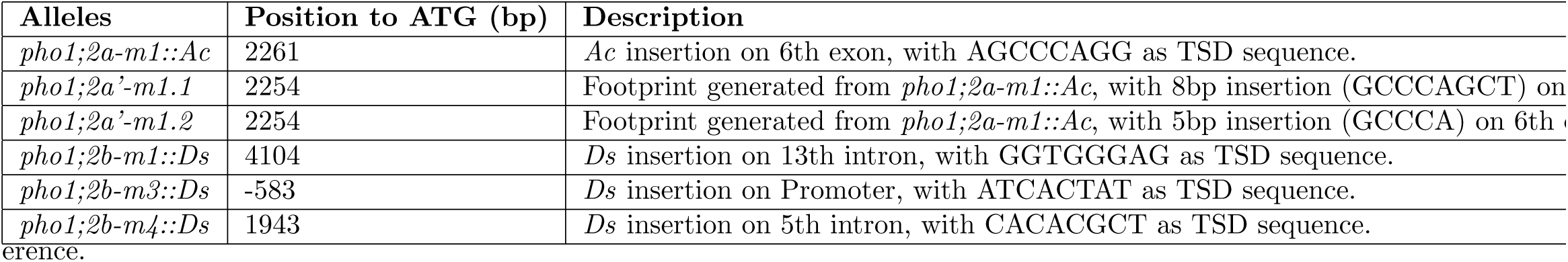
Transposon insertion alleles of maize *Pho1;2* genes

**Figure 4.**
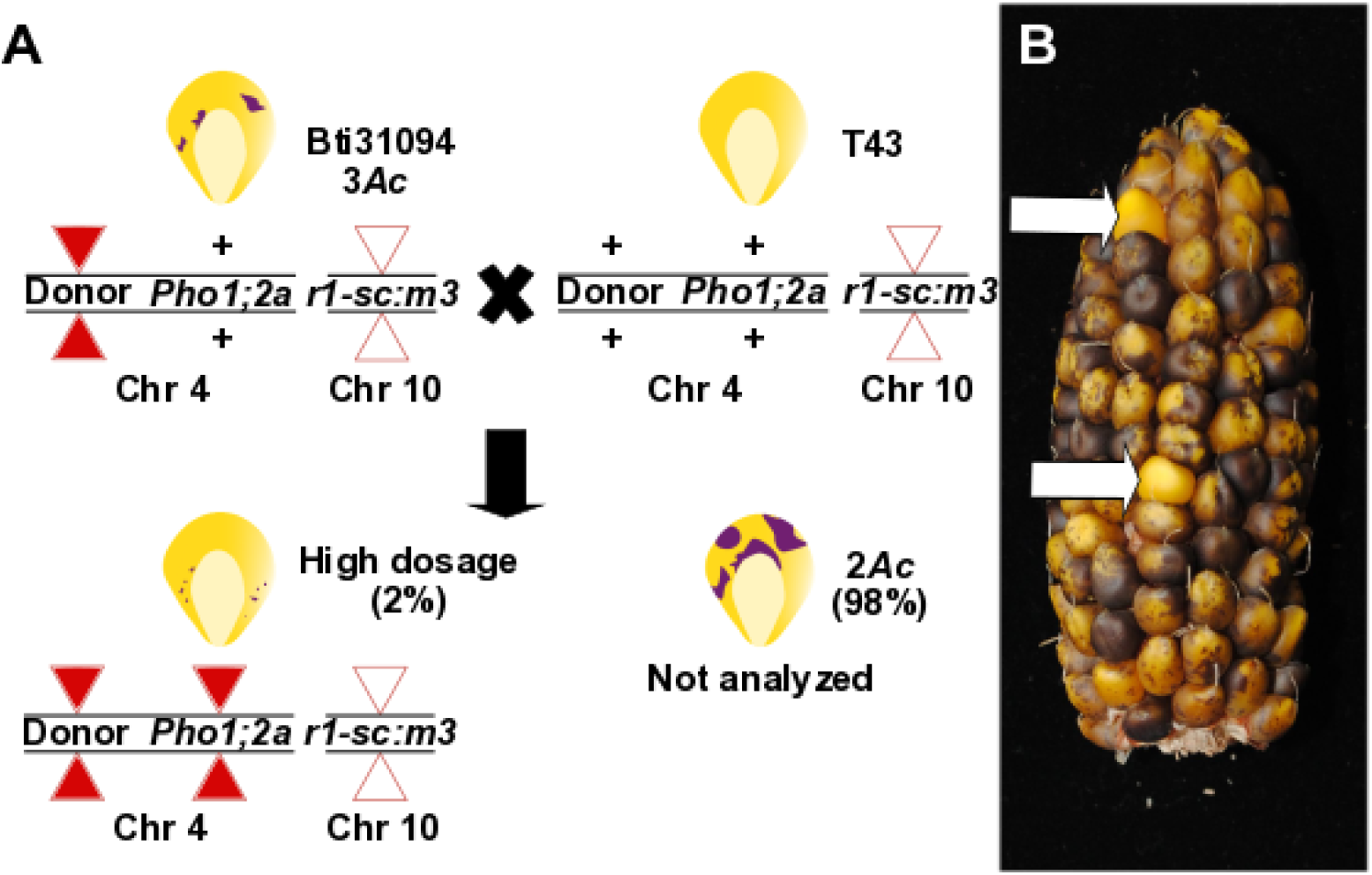
*Ac* mutagenesis of *ZmPho1;2a*. (A) Genetic strategy to mobilize the donor element *bti31094*∷*Ac.* Individuals homozygous for the donor element, displaying the characteristic 3 *Ac* dosage pattern of excision from *r1-sc:m3* in the triploid aleurone, were crossed as females by T43. A small proportion (^∼^2%) of the progeny kernels were finely spotted, indicating high *Ac* dose as a result of replicative transposition, and were screened for potential re-insertion into the target gene. (B) Two high *Ac* dose kernels (white arrows) highlighted among the largely 2 *Ac* dose progeny of a typical test cross ear.

**Figure 5.**
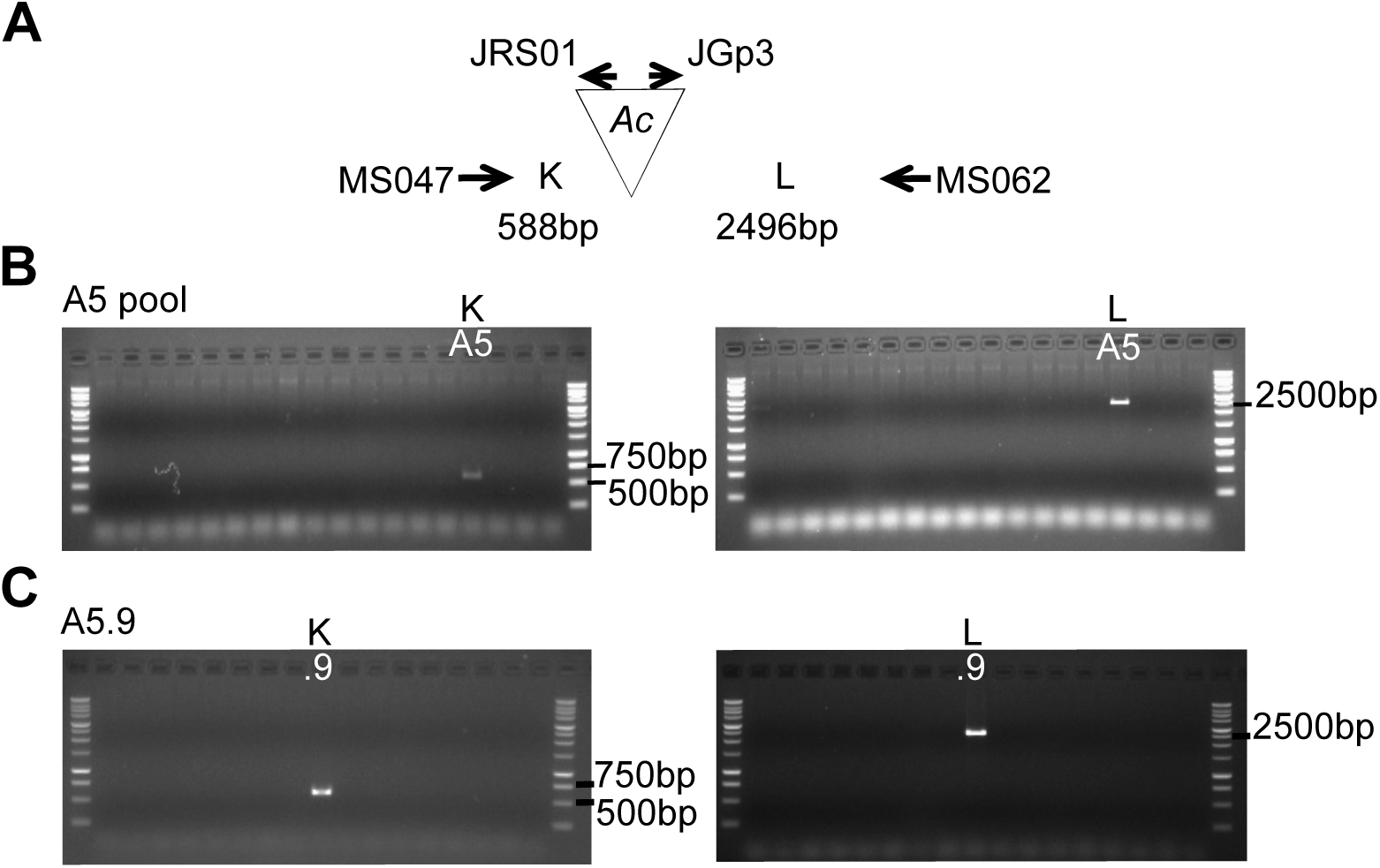
PCR screening for novel *Ac* insertion in ZmPho1;2a. (A) Diagram of expected fragment size on A5.9 allele. Reaction K (588bp) and reaction L (2496bp) are indicated. (B) PCR reactions using gene and *Ac* specific primers to amplify potential products from DNA pools. Pool A5 shows amplification in complementary reactions K and L. (C) PCR reactions using individual DNA. Detection of strong amplification from the individual A5.9 in both complementary reactions.

**Figure 6.**
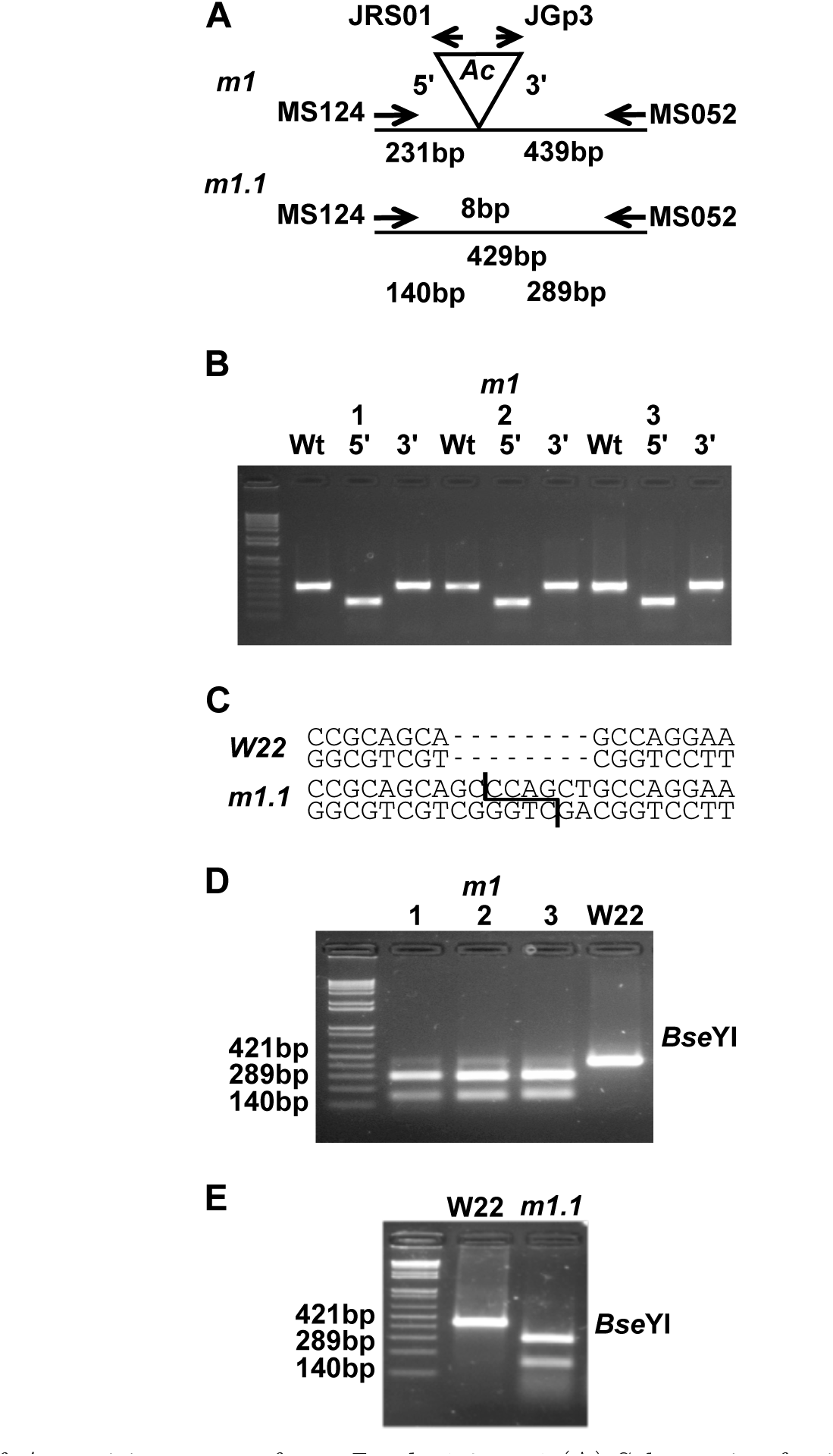
Recovery of Ac excision events from *ZmPho1;2a-m1* (A) Schematic of primer position and fragment lengths corresponding to *ZmPho1;2a-m1* (*m1*) and derived excision events, e.g. *ZmPho1;2a’m1.1 (m1.1)*. (B) Amplification of apparent wild-type (Wt) and flanking products (5’, 3’) from three different individuals selected as homozygous *ZmPho1;2a-m1* on the basis of kernal phenotype and pedigree. (C) Generation of a BseYI cutting site, not present in wild-type (*W22*), as the result of an 8bp duplication retained following excision from *ZmPho1;2a-m1*, as in e.g. *ZmPho1;2a’m1.1 (m1.1)*. (D) Digestion of products amplified with the insertion spanning primers MS124/MS052 from three *ZmPho1;2a’m1.1* homozygous individuals shown in B and a wild-type (W22) individual. (E) Digestion of products amplified with the insertion spanning primers MS124/MS052 from a wild-type individual (W22) and an individual homozygous for the stable germinal excision allele *pho1;2a’m1.1.* 1Kb plus DNA ladder (Invitrogen) is loaded on the first lane in B,C and E.

To identify novel *Ds* insertions in *ZmPho1;2b, I.S06.1616∷Ds* (designated *ZmPho1;2b-m1∷Ds*), a *Ds* element identified to be inserted in intron 13 of the target gene, was re-mobilized. Plants homozygous for the *ZmPho1;2b-m1∷Ds* allele did not present any observable phenotype and RT-PCR analysis of transcript accumulation indicated such plants to accumulate correctly-spliced transcript to normal levels (data not shown). To derive further alleles, individuals homozygous for *ZmPho1;2b-s1∷Ds* and also carrying the unlinked stable transposase source *Ac-immobilized (Ac-im)* were crossed as males to T43 (Fig. 7),and test-cross progeny screened using a strategy similar to that employed in the mutagensis of *ZmPho1;2a*. Two novel *Ds* insertions were identified, one in the promoter region, 591bp upstream of the ATG (*ZmPho1;2b-m3∷Ds*) and the second in intron 5 (*Zmpho1;2b-m4∷Ds*). Sequencing of the region flanking each novel insertions identified the expected 8bp target site duplication (Table 4).

**Figure 7.**
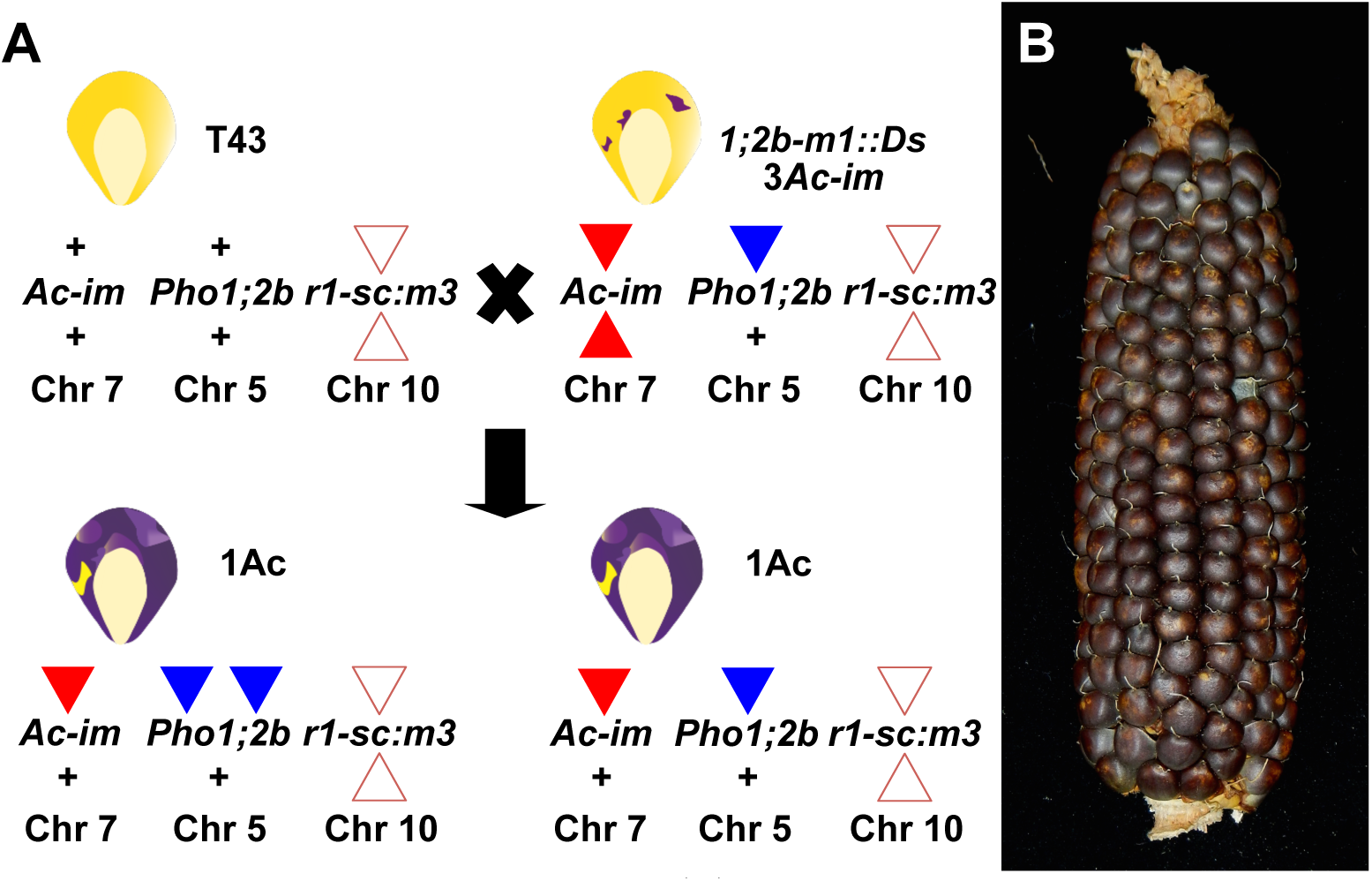
Ds mutagenesis of *ZmPho1;2b*. (A) Genetic strategy to re-mobilize the Ds element from *ZmPho1;2b-m1*∷Ds. Individuals carrying *ZmPho1;2b-m1*∷Ds and the stable transposase source *Ac-im* were crossed as males to T43. All progeny were screened as rare kernels carrying transposed Ds were indistinguishable from other progeny on the basis of aluerone phenotype. (B) Ear carrying kernels with one copy of Ac-im in the chr7 expressed in the triploid aleurone.

**Figure 8.**
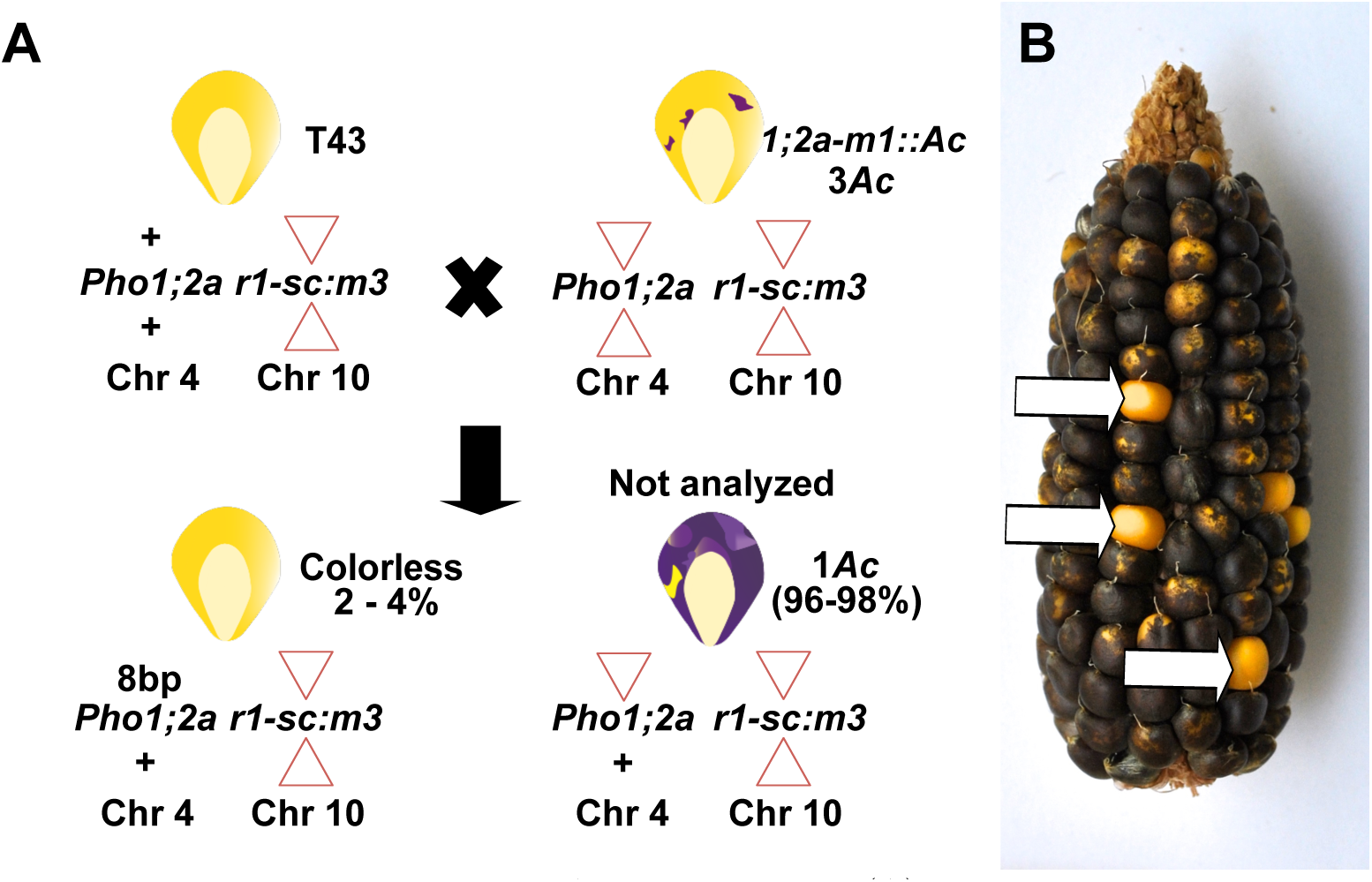
Footprint generation by Ac excision event. (A) Genetic strategy to re-mobilize to recover stable germinal excision of *Ac* from *ZmPho1;2a-m1∷Ac*. An individual homozygous for *ZmPho1;2a-m1∷Ac* was crossed as male to T43. Rare colorless kernels were selected for PCR screening. (B) Colorless kernels (white arrows) highlighted among the largely 1 *Ac* dose progeny of a typical test cross ear

### Derivation of stable derivatives of *ZmPho1;2a*

To generate stable “footprint” alleles by Ac excision from *ZmPho1;2a-m1∷Ac*, a homozygous *ZmPho1;2a-m1∷Ac* individual was crossed as a male onto T43 females, and resulting colorless progeny screened by PCR amplification across the genomic the *ZmPho1;2a-m1∷Ac* insertion site. Products of a size consistent with Ac excision were cloned and sequenced. Two footprint alleles were identified, one with an 8bp duplication (GCCCAGCT) (*ZmPho1;2a’-m1.1*) and the second with a 5bp duplication (GCCCA) (*ZmPho1;2a’-m1.2*). For confirmation, the region spanning *ZmPho1;2a’m1.1* was re-amplified and digested with the enzyme BseYI that specifically recognized a target site generated by the 8bp duplication (Fig.6A,C,E). As a result of non-triplet duplication, both *ZmPho1;2a'-m1.1* and *ZmPho1;2a'-m1.2* alleles disrupt the DNA reading frame and are predicted to result in a premature termination of translation (Table 4).

## Discussion

Maize is the most widely grown cereal in the world (ref http://faostat3.fao.org). Much of this cultivated area is P limited. And yet, the molecular basis of P uptake and translocation in maize remains poorly characterized (reviewed in [10]). In this study, we have described the maize *Pho1* gene family and generated novel mutant insertion alleles of *ZmPho1;2* genes using the endogenous maize *Ac/Ds* transposon system. The genetic material described here initiates the functional analysis of P homeostasis in maize.

The maize *Pho1* family consists of four genes, corresponding to the three gene (*PHO1;1, PHO1;2, PHO1;3*) structure reported previously in rice (*Oryza sativa*) [5], with the elaboration of the duplication of *PHO1;2*. The sorghum Pho1 family was also found to consist of three genes. The restricted PHO1 family present in these cereals is in contrast to larger 11-member family of *Arabidopsis* [9]. Specifically, the cereals lack a large clade of *PHO1* related sequences present in *Arabidopsis* that has been implicated in a range of biological functions extending beyond P homeostasis [9,20,22]. Indeed, in experiments to complement the *Atpho1* phenotype by expression of other *Arabidopsis PHO1* family members, it was only *AtPHO1;H1* that could rescue the mutant [20]. Although phylogenetic analysis and experimental data from *Arabidopsis* and rice suggest all four maize PHO1 to be directly involved in P homeostasis, further work in heterologous systems, and ultimately the analysis of the mutants described here, will be required to determine functional equivalence across species and the biological role in maize.

The lineage leading to maize experienced a tetraploidy event resulting in whole genome duplication (WGD) sometime after the split with the sorghum lineage, 5-12 million years ago [21]. Considering contemporary sorghum to represent the pre-duplication state, and taking the structure of the rice *PHO1* family into account, immediately following the tetraploid event, maize would have carried six Pho1 genes, represented by three pairs of syntenic paralogs (homeologs). Subsequently, the maize genome has returned to a diploid state through a process of reorganization that has been coupled with extensive fractionation - the loss of one of a pair of syntenic paralogs [23]. Large scale gene loss following WGD appears to be a general trend observed across taxa and across timescales [24]. Gene loss is presumed to be buffered by the presence of a functionally equivalent paralogs. Where both paralogs of a syntenic pair are retained, it may indicate either selection or simply incomplete fractionation. The former case would imply functional divergence or a selective advantag ^∼^20% of the total complement of ^∼^32,000 total genes, or closer to ^∼^10% of the pre-duplication gene set [21] [25]. While gene loss is the more likely outcome following genome duplication, it is difficult to determine the balance of selective gene-by-gene reduction and the largely random loss of larger sections of DNA. Similarly, where a pair of syntenic paralogs are retained, as is the case with *Pho1;2*, it may indicate selection directly on the gene pair or a genomic context that insulates the gene pair from larger scale DNA loss events. It is noticeable that a number of syntenic paralog pairs have been retained close to the *Pho1;2* locus, potentially “hitchhiking” on direct selection to maintain one or more of the adjacent pairs. In the case of the pair GRMZM2G164854/GRMZM5G853379, the two paralogs overlap directly with *Pho1;2* sequence on the opposite DNA strand. Consequently, selection to maintain either the *Pho1;2* or GRMZM2G164854/GRMZM5G853379 paralog pair might protect also the adjacent genes from silencing or deletion.

Analysis of *ZmPho1;2a* and *ZmPho1;2b* transcript accumulation demonstrated regulatory divergence between the two paralogs, with *ZmPho1;2b* transcripts presenting a pattern more similar to that of *SbPho1;2*, the presumed pre-duplication state. Characterization of the leaf transcriptome has estimated 13% of retained syntenic paralogs to undergo regulatory neo-functionalization [25], placing the *Pho1;2* pair among just 2-3% of the total maize gene set. Characterization of putative *cis*-NAT transcripts offered further evidence of regulatory divergence between maize Pho1;2 paralogs. Accumulation of *cis*-NAT_*ZmPho1;2a*_ was induced by P limitation, in a manner similar to that observed for *cis*-NAT_*OsPho1;2*_, while *cis*-NAT_*ZmPho1;2b*_ accumulation mirrored that of the *ZmPho1;2b* sense transcript. Given a failure to detect *Pho1;2* associated anti-sense transcripts in sorghum using the techniques applied, we might infer the pre-duplication state from the more distantly related rice. Interestingly, on such a basis, our data are consistent with regulatory neo-functionalization acting on the one hand on *ZmPho1;2a* sense, and on the other on *ZmPho1;2b* anti-sense, transcript accumulation. Although characterized *cis*-NATs act on the activity of adjacent protein coding genes, the translational enhancer function postulated for *PHO1;2* NATs may allow for trans action between *ZmPho1;2* paralogs given the degree of sequence similarity. Indeed, one intriguing hypothesis, suggested by our transcript accumulation data, is that the situation might be best considered as sub-functionalization of maize *PHO1;2*, with the primary production of sense transcripts from *ZmPho1;2b* and the primary production of anti-sense transcripts from *ZmPho1;2a*.

Characterization of the insertional alleles described here will be central in determining the function of *ZmPho1;2a* and *ZmPho1;2b*. We are continuing to mobilize Ac and Ds elements at the maize *Pho1;2* loci, taking full advantage of the capacity of the system to generate allelic series, impacting variously sense and anti-sense transcripts. Such material will be invaluable in the fine-scale evaluation of regulatory crosstalk and functional redundancy between between *ZmPho1;2* paralogs and, ultimately, the biological role of PHO1 proteins in maize.

## Acknowledgments

We thank Juan Estevez-Palmas for valuable comments on the manuscript. This work was supported by the National Science Foundation grant IOS-0922701 to TB and the Mexican National Council of Science and Technology (CONACYT) grant CB2012-151947 to RS.

